# Absorbing molecules make both abdomen and back transparent in live mice

**DOI:** 10.1101/2024.10.28.620537

**Authors:** Xin Tie, Ting Sun, Guixiu Xiao, Yanjie Zhao, Jing Su, Xiaoqi Xie, Wanhong Yin

## Abstract

The field of tissue clearing has experienced rapid advancements in the past decades. Tissue clearing techniques primarily rely on physical or chemical approaches to match the refractive indices of cells and their surrounding medium, thereby reducing light scattering and rendering the tissue optically transparent. By doing so, it becomes possible to utilize fluorescence imaging and 3D reconstruction techniques to observe and analyze the contents and structures of tissues at a cellular level. However, effective research on in vivo tissue clearing remains limited due to the involvement of toxic substances and the removal of water or lipids. A recent article in *Science* reports a technique that achieves tissue transparency in live animals with absorbing molecules, marking a significant advancement in the field of in vivo tissue clearing. Although this work has great potential for imaging and light-based therapeutics, some labs found it difficult to reproduce the reported results. In this study, we report the successful reproduction of the findings, provide experimental details that we identified as key to reproducibility, present a new transparency effect observed in the back of live mice, which was not shown in the original paper, and offer insights into the future development and applications of in vivo clearing reagents.

## Introduction

Tissue clearing technology has expanded traditional 2D histological sectioning into 3D reconstruction, allowing for the preservation of more intricate biological information [1] [2]. This enables researchers to delve deeper into the biological mechanisms underlying health and disease. The technique achieves optical transparency by matching the refractive indices of different tissue components, such as lipids, proteins, and water, thereby reducing light scattering [3] [4] [5]. However, current tissue-clearing methods are primarily applied to post-mortem animal tissues, and the use of organic solvents or other chemicals during processing may result in the loss of key tissue components and biological information. Additionally, tissue transparency achieved during the clearing process often causes tissue shrinkage or expansion due to dehydration or hyperhydration [6]. Moreover, fluorescence signals may weaken, and staining can become inconsistent during subsequent labeling or staining steps, potentially introducing errors in the analysis [7].

Recently, Dr. Hong’s team developed a novel approach that uses the edible dye tartrazine to achieve transparency in live mice, enabling macroscopic visualization of abdominal organs and their movements. The technique also allows microscopic observation of details in the hindlimb muscles and intracranial vasculature [8]. After reviewing Dr. Hong’s findings, we became highly interested in this research and attempted to reproduce their results. In this work, we report our replication of the reported results, providing important details to facilitate reproducibility based on our own experiments, along with new findings showing transparency in the back of live mice. We aim to further explore potential applications of this technique in future medical fields.

## Materials and Methods

### Chemicals

We used tartrazine (T0388, industrial grade, ≥85%) and 4-aminoantipyrine (4-AA, A4382, reagent grade) sourced from Sigma Aldrich, along with low-melting-point agarose (A9414, CAS number: 39346-81-1), also purchased from Sigma Aldrich.

### Animals

Female C57BL/6 mice and Sprague-Dawley (SD) rats were obtained from BEIJING HFK BIOSCIENCE CO, LTD (Beijing, China). These animals were housed under specific pathogen-free conditions and treated humanely throughout the study. Ex vivo tissue samples, after dissection, dorsal skin samples from SD rats were promptly collected and sectioned into approximately 1 cm x 1 cm squares for experimental use. Ex vivo tissue samples were immersed in tartrazine solutions with concentrations of 20 mM, 163 mM, 0.62 M, and 0 M, respectively, for 1 day, and tissue transparency was then observed (**Figure 2**).

### Gel preparation for in vivo transparency

We weighed out tartrazine and 4-aminoantipyrine (Detailed amounts are specified in **Table 1**, according to the *Science* paper [8]) and transferred them into a 20 mL reagent bottle. We then added distilled water to the bottle, ensuring thorough mixing by vortexing. We placed the bottle in an oven preheated to 70-80°C and let it sit for 10 min to ensure complete dissolution of the tartrazine or 4-aminoantipyrine. Then, we added 30 mg of low-melting-point agarose to the fully dissolved solution. We returned the bottle to the 70-80°C oven and let it sit for another 10 minutes to allow the agarose to dissolve completely. We then gently swirled the bottle to ensure the agarose was evenly distributed throughout the solution, resulting in a homogeneous mixture (**Figure 1**).

**Table 1.**
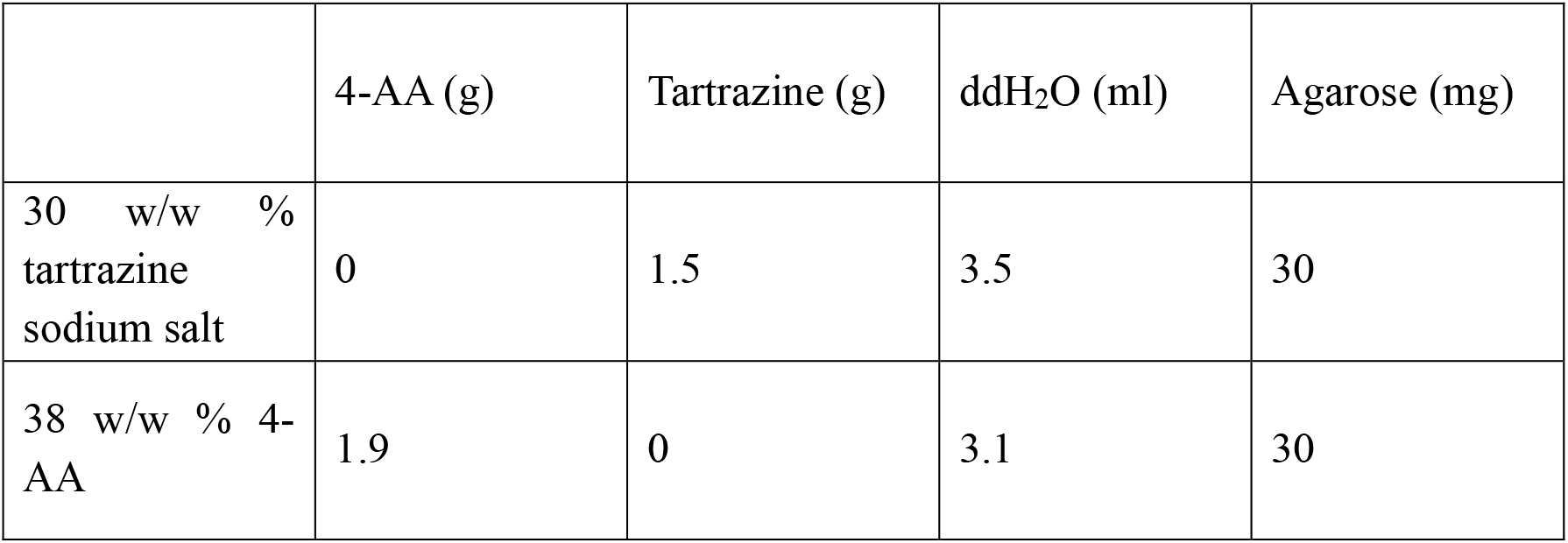
Composition of tartrazine and 4-AA gels.

**Figure 1.**
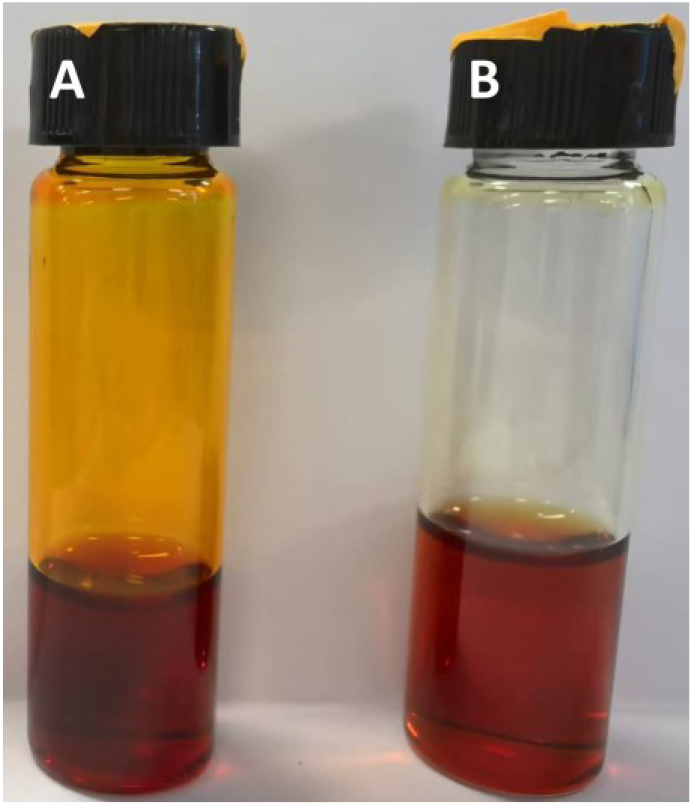
Tartrazine and 4-AA solutions. (A) 30 w/w% tartrazine solution; (B) 38 w/w% 4-AA solution.

**Figure 2.**
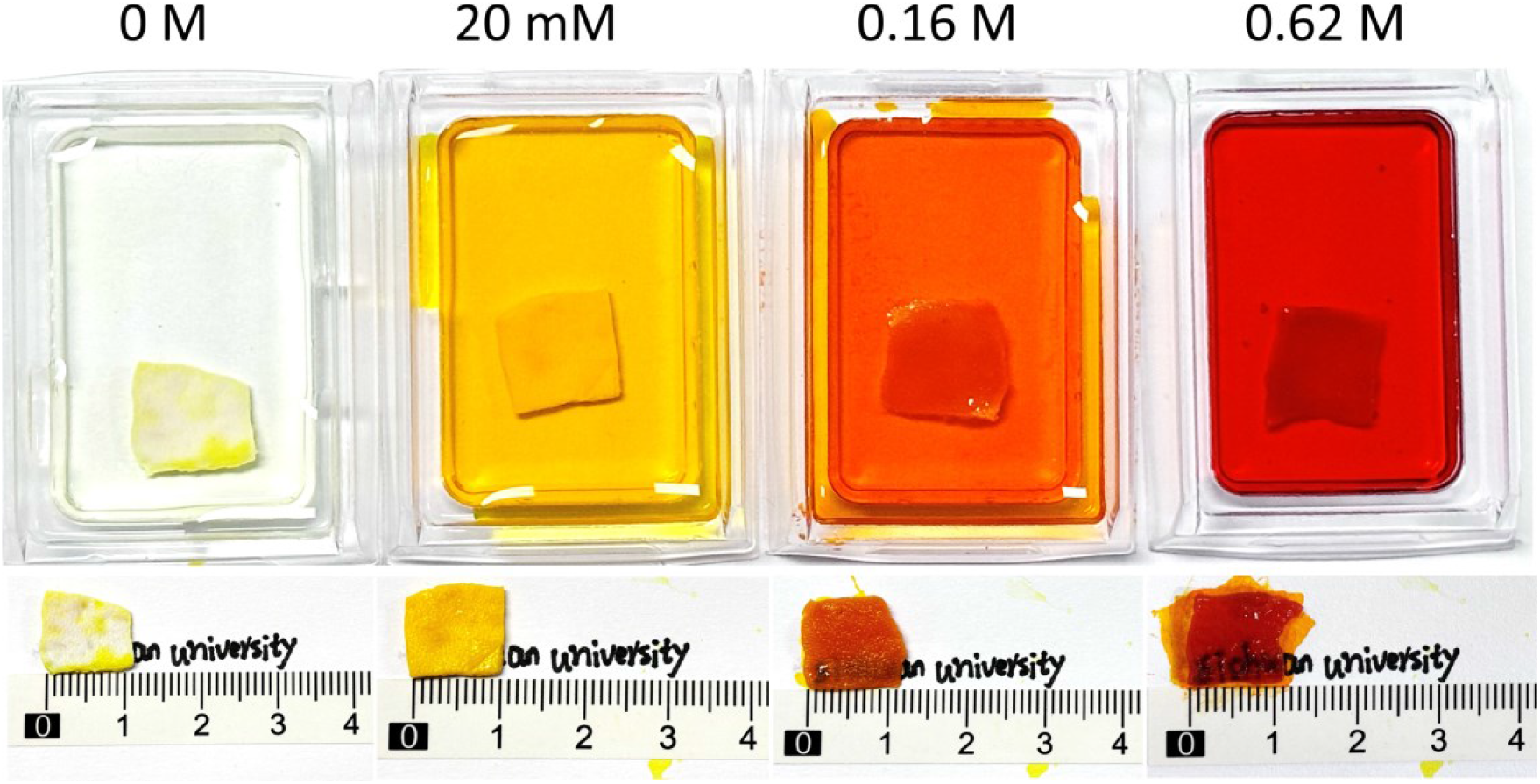
Rat skin soaked at 0 M, 20 mM, 163 mM, and 0.62 M concentrations of tartrazine solution. Labels on the ruler are in centimeters.

### Animal experiments

We selected 3-to 4-week-old C57BL/6 mice weighing between 8 and 15 grams. We anesthetized the animals using inhalation or intraperitoneal injection. Depilatory cream was used to first remove all hair on the abdomen. We then applied depilatory cream again generously to the abdominal area intended for clearing and waited briefly before removing it thoroughly with an alcohol wipe. We then applied the gel containing the absorbing molecules to the desired area using a cotton swab, rubbing gently. After massaging the gel onto the skin for 5 minutes, the treated area turned from orange to red, with increased translucency. We continued massaging for an additional 5 minutes to achieve maximal transparency. For mice treated with 4-AA solution, skin translucency began to appear within 1 minute of massaging, reaching maximum transparency and window clarity after a total of 5 minutes. For photography of the transparent window with minimal glare, we applied one drop of glycerol on the cleared region and used a glass slide to cover the area of interest. To reverse skin transparency, we thoroughly rinsed the area with warm water or saline to remove any residual dye. After rinsing, the skin returned to its normal opacity. To prevent skin inflammation, we applied a thin layer of petroleum jelly ointment to the treated area. Upon recovery from anesthesia, the animal returned to its normal movement while we monitored the treated area and reapplied ointment as needed.

## Results and discussion

In the ex vivo experiments, we used relatively thick rat skin samples (2 mm, **Figure 2**), while for the in vivo studies, we utilized smaller C57BL/6 mice (3-4 weeks old), which have significantly thinner skin. This discrepancy in skin thickness necessitated different application times and clearing durations, with potentially longer application times required for in vivo SD rats. Consequently, it is reasonable to infer that the observed transparency window may be smaller in live SD rats, adding to the challenges in applying this technique.

In the in vivo experiments, both tartrazine and 4-aminoantipyrine exhibited abdominal transparency (**Figure 3** and **Figure 4**). Additionally, we observed that 4-aminoantipyrine exhibited superior clearing effects compared to tartrazine, with a faster onset of transparency (visible within 1 minute, **Figure 4B**) and much less red color in the cleared window. We attributed this difference to the smaller molecular weight of 4-AA and its faster diffusion through the skin. Therefore, future improvements to tartrazine-based reagents should focus on enhancing permeability and prolonging the duration of the transparency effect while maintaining reagent safety. Lastly, by applying the tartrazine gel to the back of the mice, we observed the spleen and the spine through the cleared skin and subcutaneous tissues in live mice (**Figure 4A**). This is a new observation not reported in the original *Science* paper and suggests broader applications of this technique.

**Figure 3.**
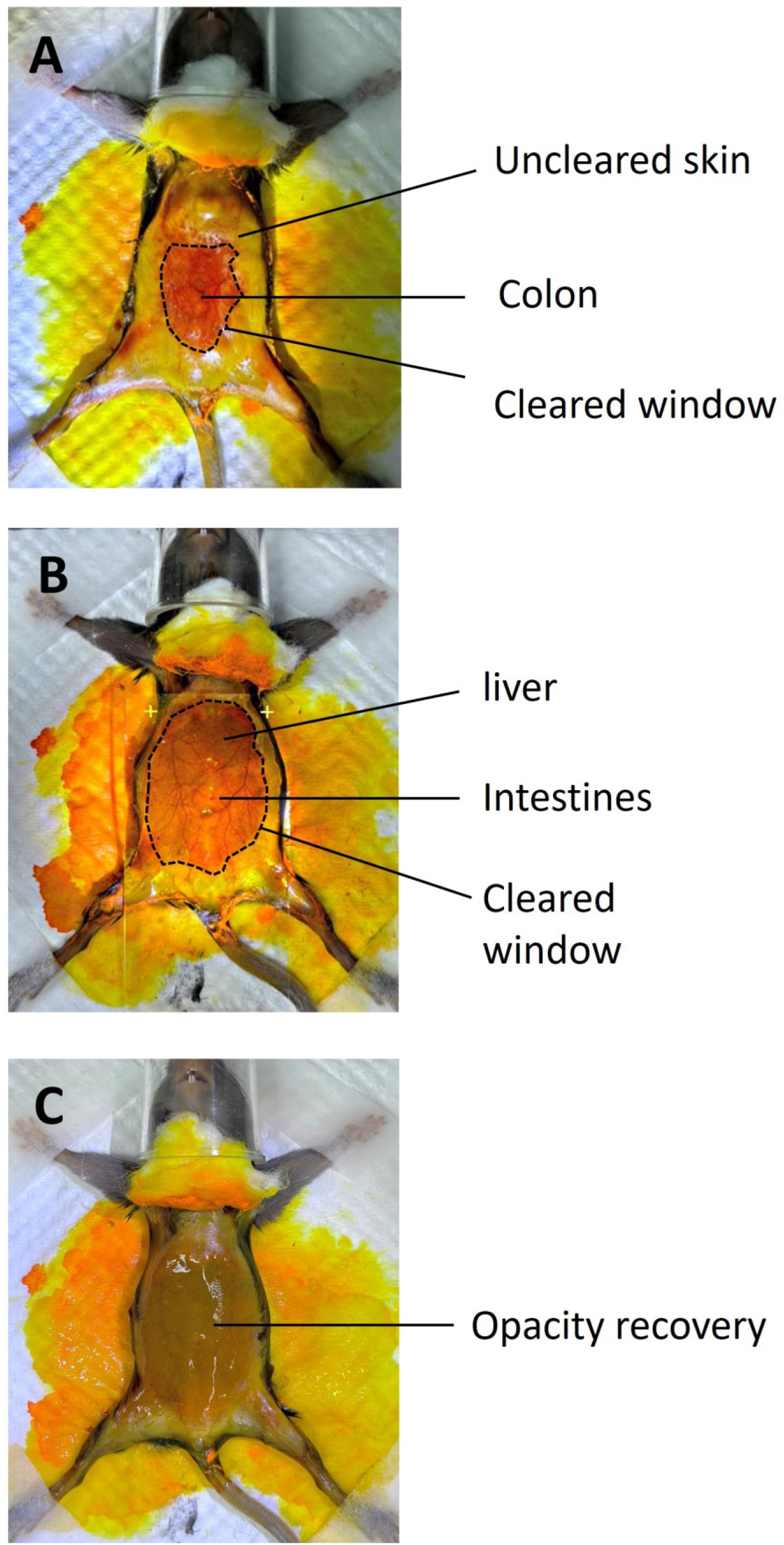
In mice treated with tartrazine gel, transparency began at 5 min (A), maximum transparency and window were reached at 10 min (B), and the degree of transparency gradually disappeared after scrubbing the skin with water (C).

**Figure 4.**
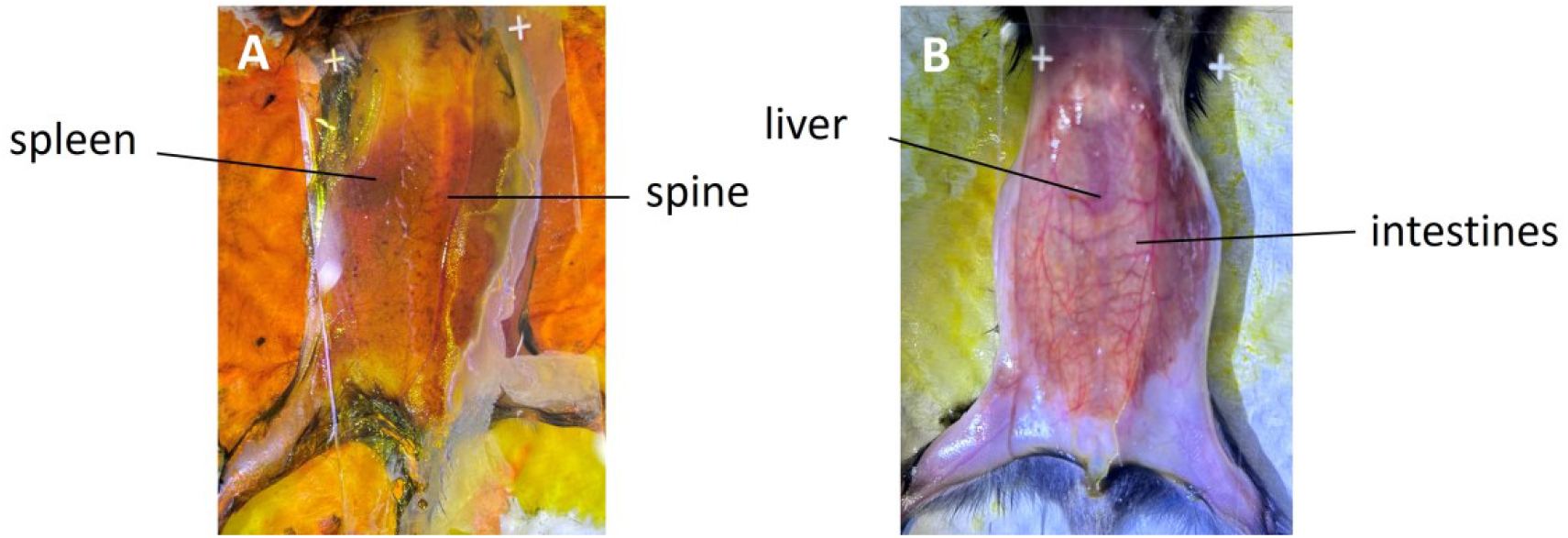
(A) The spleen and spine became visible after applying the tartrazine gel to the back of the mice. (B) 4-AA gel was applied to the abdomen of mice, and the liver and colon became visible after application with much less red color than tartrazine.

As shown by our experimental results, we successfully replicated the findings reported in *Science*. In our live mouse experiments, we observed the morphology and activity of internal organs, such as the liver and intestines, through a transparency window created by skin clearing. We also conducted experimental applications in other scenarios, such as spinal observation, and demonstrated that even denser, thicker rat skin tissues can become transparent using the same solution. These findings strongly validate the feasibility and future scalability of in vivo skin clearing technology.

This in vivo tissue clearing method could enable detailed observations of biological tissues without compromising structural integrity, opening new avenues for both basic and clinical research [6] [9]. As the largest organ of the human body, the skin not only protects internal organs but also plays a crucial role in disease diagnosis and treatment guidance. However, the inability to view deeper tissues through the skin often necessitates invasive or radiological examinations for diagnosing and treating many conditions. By applying tissue clearing to the skin, we can obtain an unbiased, system-level view of human bodies and specimens that aids in diagnosing, researching, and preventing various diseases.

Our experimental results suggest broad future applications for skin transparency technology. Subcutaneous masses, which manifest as lumps, nodules, or swellings in deep skin layers and arise from various causes, could be diagnosed more effectively using this technology. It could aid in distinguishing these masses from malignant tumors, such as skin cancer [10]. Moreover, transparency enhances the visibility of subcutaneous veins and arteries, significantly increasing the success rate of procedures like venipuncture and injections, thus reducing patient discomfort and infection risks [11]. In the cosmetic field, these reagents could help mask skin imperfections such as pigmentation and wrinkles [12].

Observing intestinal motion in vivo through transparency leads us to consider pediatric complications like intussusception and life-threatening conditions such as congenital megacolon [13] [14]. Establishing a transparent observation window in the abdomen could allow straightforward visualization of intestinal abnormalities, potentially reducing CT false-positive rates and minimizing medically induced radiation exposure in infants. Abdominal effusions, excluding those in deeper tissue spaces, could also be observed directly [12]. For cataract patients, where protein denaturation in the lens severely impairs vision, the introduction of tartrazine-like agents into the lens for clearing may offer new hope for vision restoration [15].

In vivo skin transparency technology also shows vast potential for future applications in microvascular observation. Microcirculatory dysfunction is a common issue among ICU patients, and early detection of affected microcirculatory sites can significantly reduce mortality rates and alleviate healthcare resource demands [16]. Diabetic microangiopathy often first appears in the lower extremities, particularly in the microvasculature of the anterior tibia and foot skin. This condition is characterized by capillary basement membrane thickening, which impairs blood flow, leading to tissue hypoxia and nutrient deficiency, and eventually resulting in ulceration. Early observation of microvascular abnormalities in diabetic patients and timely blood glucose control could greatly reduce the risk of diabetic foot ulcers [17].

Most importantly, once the safety and efficacy of tartrazine, 4-aminoantipyrine, and other absorbing molecules are thoroughly validated, they may hold promise for use in maternity care, facilitating the early detection of fetal malformations and uterine abnormalities. We believe that the creation of transparent observation windows on the surface of live animals could have a profound impact on the progression of various diseases.

In summary, in vivo tissue clearing technology holds tremendous potential for both basic research and clinical applications. By facilitating the clearing of skin and other tissues, this technique not only allows for in-depth observation of the structure and dynamics of various organs within the body but also plays a unique role in early disease diagnosis, therapeutic assistance, and cosmetic care. With ongoing improvements in the safety and efficacy of reagents such as tartrazine and 4-aminoantipyrine, in vivo tissue clearing technology is expected to be further applied in human medical diagnostics, potentially aiding in the early detection and intervention of complex diseases. In the future, the integration of additional non-invasive imaging techniques and the optimization of the permeability and imaging effects of clearing agents could lead to innovative developments in the field of biomedicine.

## Material availability statement

Stable solutions and gels containing tartrazine and 4-AA can be distributed to interested labs on request. Requests to these experimental materials should be directed to.

## Author contribution

Xin Tie and Ting Sun jointly designed and conducted the experiments and completed the manuscript writing. Guixiu Xiao participated in the experiments. Yanjie Zhao and Jing Su contributed to manuscript writing. Xiaoqi Xie and Wanhong Yin provided support for the project and offered revision suggestions.

